# Combined effect of baicalein and thermal-cycling stimulation on suppressing non-small cell lung cancer A549 cells under CoCl_2_-induced hypoxia

**DOI:** 10.64898/2026.08.11.744169

**Authors:** Yu-Wei Wang, Guan-Bo Lin, Fang-Tzu Hsu, Yu-Yi Kuo, Yun-Hao Chen, Chih-Yu Chao

## Abstract

Lung cancer continues to be the leading cause of cancer-related mortality globally, with non-small cell lung cancer (NSCLC) representing the most prevalent subtype. Tumor hypoxia is a characteristic feature of the neoplastic microenvironment in NSCLC, facilitating tumor progression and conferring resistance to oxidative stress through the stabilization of hypoxia-inducible factor-1 alpha (HIF-1α). In this study, we investigated the combined anticancer effects of baicalein (Bai), a natural flavonoid, and thermal-cycling stimulation (TCS), a physical treatment that minimizes damage to normal cells, under cobalt (II) chloride (CoCl_2_)-induced hypoxic conditions in NSCLC. In A549 NSCLC cells, the combination of Bai and TCS significantly decreased cell viability and induced apoptosis, while exhibiting minimal cytotoxicity on IMR-90 normal human lung fibroblast cells. On a mechanistic level, this combined treatment suppressed the expression of HIF-1α and superoxide dismutase 2 (SOD2) proteins, elevated intracellular reactive oxygen species (ROS) levels, and impaired DNA repair capability by downregulating MutT homolog 1 (MTH1) protein expression. Additionally, disruption of mitochondrial membrane potential and increased poly (ADP-ribose) polymerase (PARP) cleavage further confirmed the induction of apoptosis. These findings indicate that combining Bai with TCS offers a promising synergistic approach to treating NSCLC under hypoxic conditions.

## Introduction

In recent years, lung cancer has surpassed colorectal cancer in both incidence and mortality, making it the leading cause of cancer-related deaths [1]. As a result, the treatment and research of lung cancer have become increasingly important. Non-small cell lung cancer (NSCLC) is the most common subtype, comprising about 80–85% of all lung cancer cases [2], which highlights the critical need for the development of effective therapeutic interventions specifically targeting this disease subtype. Although current treatments such as surgery, radiotherapy, and chemotherapy are commonly used for lung cancer [3], they often come with significant side effects [4, 5]. Therefore, it is crucial to develop new therapeutic strategies that can reduce adverse effects while maintaining anticancer effectiveness.

A hypoxic microenvironment is commonly observed in solid tumors, mainly because abnormal and disorganized vasculature cannot meet the oxygen demands of rapidly proliferating tumor cells. This leads to limited oxygen diffusion while the metabolic demand continues to increase, resulting in a persistent hypoxic state [6, 7]. In such conditions, cells stabilize and activate hypoxia-inducible factors (HIFs), a group of transcription factors that regulate cellular responses to low oxygen levels [7, 8]. Among the HIF family, HIF-1 is the most extensively characterized and functions as a heterodimer comprising of HIF-1α and HIF-1β subunits [9]. Under normoxic conditions, HIF-1α undergoes rapid degradation via oxygen-dependent post-translational modifications. In contrast, hypoxic conditions inhibit this degradation pathway, leading to the stabilization of HIF-1α and subsequent activation of genes that facilitate cell survival, metabolic adaptation, and disease progression [10–13]. *In vitro*, cobalt (II) chloride (CoCl_2_) is widely used as a chemical hypoxia-mimetic agent due to its efficacy in stabilizing HIF-1α and thereby inducing hypoxia-related signaling pathways [14]. Elevated expression of HIF-1α is a recognized feature of hypoxic tumors and is closely associated with increased cellular proliferation and resistance to cellular stress [10–13]. A critical downstream effect of HIF-1α activation involves modulation of the cellular redox balance. Under hypoxic conditions, reactive oxygen species (ROS) are produced as by-products of aerobic metabolism and cell growth [15, 16]. When ROS levels become too high, they overwhelm antioxidant defenses, causing oxidative stress and cellular damage [17]. To mitigate this, HIF-1α has been shown to increase the production of the mitochondrial antioxidant enzyme superoxide dismutase 2 (SOD2) [18], which converts harmful superoxide radicals into hydrogen peroxide (H_2_O_2_) [19], thereby maintaining redox homeostasis and protecting cells from oxidative damage. This adaptive response may enhance the capacity of cancer cells to withstand oxidative stress in hypoxic environments [15, 18]. Given HIF-1α’s crucial role in these processes, therapeutic approaches that concurrently promote intracellular ROS generation and disrupt HIF-1α signaling in tumor cells may provide significant advantages in the treatment of NSCLC.

Baicalein (Bai) is a natural flavonoid derived from *Scutellaria baicalensis*, a traditional medicinal herb in the Lamiaceae family [20]. Previous studies have demonstrated that Bai can inhibit the HIF-1α expression in gastric cancer cells, esophageal cancer cells, and orthotopic glioma models [21–23]. Nevertheless, whether Bai similarly regulates HIF-1α expression in NSCLC cells under hypoxic conditions remains unclear. In the research of lung cancer, Bai has been evaluated using the A549 NSCLC cell line, where it has been associated with multiple anticancer mechanisms [24–26]. For instance, Bai has been shown to increase intracellular ROS levels and disrupt mitochondrial membrane potential (MMP), thereby inducing apoptosis in A549 cells [24]. These findings indicate that Bai may possess considerable therapeutic potential as a treatment for NSCLC. Despite its potential, Bai is hindered by significant pharmacokinetic challenges, including poor aqueous solubility and extensive first-pass metabolism in the intestine and liver of both humans and rodents [25, 26], which greatly limits its systemic bioavailability. Indeed, the absolute oral bioavailability of Bai has been reported to be approximately 36.1% [27]. Consequently, relatively high dosages are often necessary to achieve therapeutic effects. Therefore, improving the efficacy or potency of Bai is a crucial area for future research to enhance its therapeutic value in NSCLC.

In addition to pharmacological interventions, physical stimulation has emerged as a complementary modality in cancer treatment. Hyperthermia stimulation (HTS), which involves the application of localized heat to tumor tissues at temperatures between 40 and 45 °C, has been recognized as an effective method for inducing damage in cancer cells [28–30]. In A549 NSCLC cells, HTS has been demonstrated to elevate ROS production, disrupt mitochondrial MMP, and trigger apoptotic pathways [31, 32]. However, prolonged exposure to elevated temperatures poses the risk of harming adjacent healthy tissues [33]. To address this limitation, our research group previously developed thermal-cycling stimulation (TCS) as an alternative to conventional HTS [34]. TCS utilizes repeated cycles of heating and cooling, thereby permitting normal cells to avoid damage during intermittent high and low temperature changes and reducing thermal toxicity while maintaining the enhanced heat sensitivity of cancer cells [35]. Prior investigations have indicated that the combination of TCS with specific natural compounds potentiates anticancer effects in pancreatic cancer cells [34, 36]. Moreover, in NSCLC, the integration of TCS with the targeted therapeutic agent erlotinib has been reported to reduce A549 cell viability, enhance treatment effectiveness, and minimize damage to normal cells [37]. Although TCS has been proposed as a safer alternative to traditional HTS, its effects under hypoxic conditions—particularly concerning the regulation of HIF-1α—remain to be elucidated. If Bai, when used alongside with TCS, exhibits synergistic anticancer effects in hypoxic NSCLC cells and enables dose reduction, this combined approach could offer a promising therapeutic strategy.

In this study, we used A549 cells, a commonly used model for NSCLC, to investigate the potential anticancer effects of Bai combined with TCS in a CoCl_2_-induced hypoxic environment. We aimed to determine whether these two treatments could synergistically enhance anticancer efficacy while minimizing cytotoxicity to normal lung cells. Specifically, we evaluated cell viability, the expression of hypoxia-related proteins, oxidative stress levels, mitochondrial integrity, and apoptosis. The results demonstrated an elevation in ROS levels, a reduction in the expression of HIF-1α, SOD2, and MTH1 proteins, and a synergistic decrease in cell viability as determined by the MTT assay. These findings suggest that the combination of Bai and TCS could be a promising therapeutic strategy for NSCLC under hypoxic conditions.

## Materials and methods

### Cell culture

The NSCLC cell line A549 and the normal human lung fibroblast cell line IMR-90 were obtained from the Bioresource Collection and Research Center, Food Industry Research and Development Institute (Hsinchu, Taiwan). Cells were maintained in Dulbecco’s modified Eagle’s medium (DMEM) (CORNING, Manassas, VA, USA) supplemented with 10% fetal bovine serum (HyClone; Cytiva, Marlborough, MA, USA) and 1% penicillin-streptomycin (Gibco; Thermo Fisher Scientific, Inc., Waltham, MA, USA). All cultures were incubated at 37°C in a humidified incubator containing 5% CO_2_. To mimic hypoxic conditions, cells were pretreated with DMEM containing CoCl_2_ (Sigma-Aldrich; Merck KGaA, Darmstadt, Germany) for 6 h to chemically induce hypoxia [38].

### Drug preparation

Bai (Sigma-Aldrich; Merck KGaA) was dissolved in dimethyl sulfoxide (DMSO) (Sigma-Aldrich; Merck KGaA) to create a stock solution. Working solutions were prepared by diluting this stock in complete culture medium, ensuring that the final DMSO concentration did not exceed 0.5% (v/v) in all experimental conditions [39, 40]. CoCl_2_ was dissolved in deionized and distilled water (ddH_2_O) to prepare a stock solution, which was then diluted similarly in complete culture medium to achieve the desired working concentrations.

### TCS treatment

TCS treatment was conducted using a 2720 Thermal Cycler (Applied Biosystems; Thermo Fisher Scientific, Inc., Waltham, MA, USA) as shown in Fig 1A. The TCS protocol involved heating the cells to about 43°C for 3 min, followed by a short cooling period at approximately 41°C for 30 s. This heating and cooling cycle was repeated 10 times [36, 37]. Temperature changes in the culture medium during treatment were monitored using a thermocouple thermometer, and the recorded temperature profile was presented in Fig 1B.

**Fig 1.**
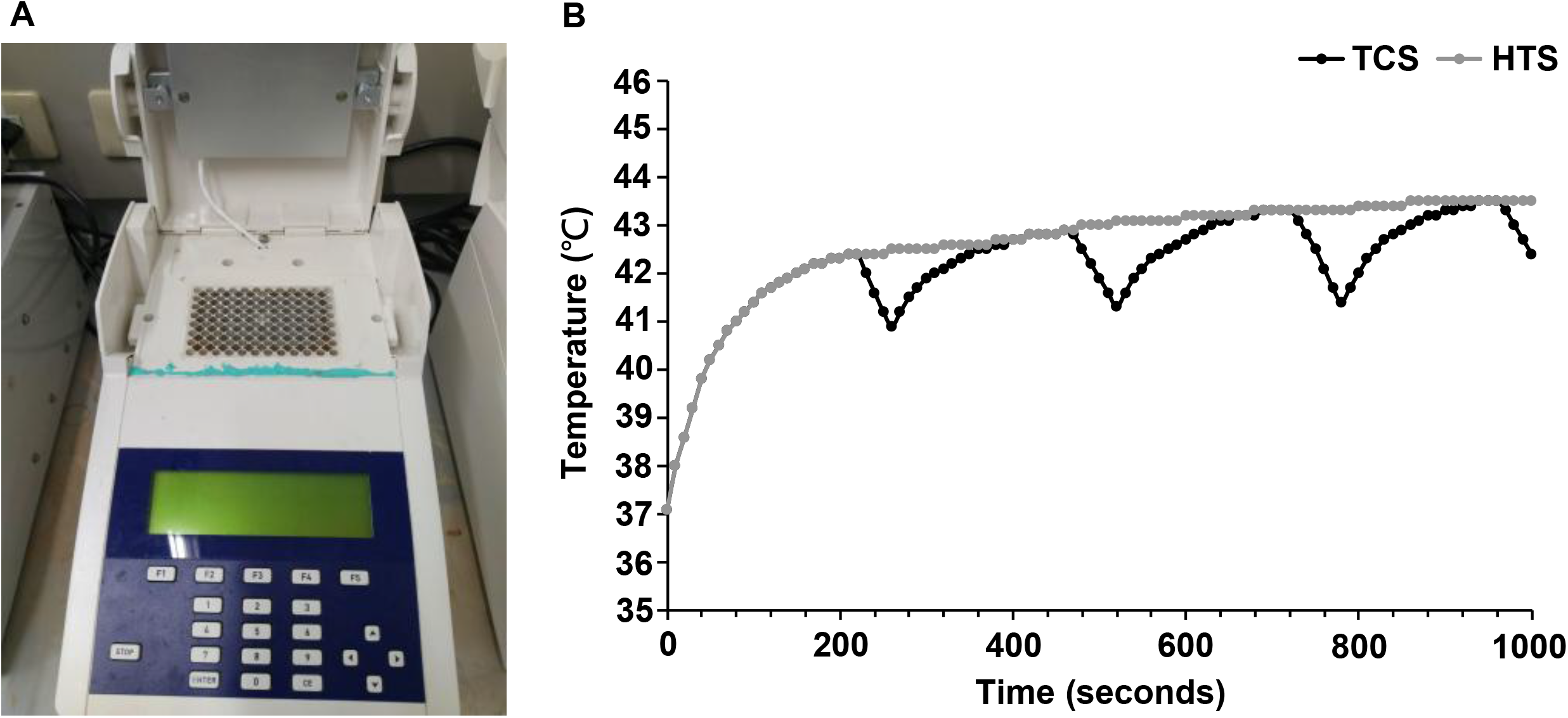
Experimental setup and temperature profiles. (A) TCS device (modified from 2720 Thermal Cycler) employed for cellular heating stimulation. (B) Temperatures in the culture medium were measured with a thermocouple thermometer during TCS and HTS treatments, illustrating the actual temperature variations experienced by the cells.

### Cell viability assay

Following Bai, TCS, or their combination treatment, A549 and IMR-90 cell viability was quantified via the MTT assay utilizing 3-(4,5-dimethylthiazol-2-yl)-2,5-diphenyltetrazolium bromide (GoldBio, St. Louis, MO, USA). The culture medium was replaced with 100 μL of MTT solution (0.5 mg/mL in DMEM) per well, and cells were incubated at 37 °C for 4 h to facilitate the enzymatic reduction of MTT to purple formazan crystals by metabolically viable cells [21, 41]. Afterwards, 100 μL of 10% sodium dodecyl sulfate (SDS) (Bioman Scientific Co., Ltd., New Taipei City, Taiwan) in 0.01 M hydrochloric acid (ECHO CHEMICAL Co., Ltd., Miaoli, Taiwan) was added to each well to solubilize the formazan precipitate. Plates were then incubated overnight at 37 °C to ensure complete dissolution. Absorbance was measured at 570 nm using a microplate reader (Thermo Fisher Scientific, Inc., Waltham, MA, USA), with background absorbance at 690 nm subtracted. The resulting optical density values were normalized to the untreated control group and expressed as relative cell viability percentages.

### Synergy quotient calculation for synergism

The synergistic effect between Bai and TCS was assessed using the synergism quotient (SQ) method as described previously [42]. Initially, the growth inhibition rates for each individual treatment were calculated. The SQ value was then determined by dividing the net inhibition rate of the combined treatment (Bai + TCS) by the sum of the inhibition rates of the individual treatments (Bai and TCS). An SQ value greater than 1.0 indicates a synergistic effect.

### Western blot analysis

Protein expression in A549 cells was examined by Western blot analysis. Following treatment with Bai alone, TCS alone, or their combination, cells were washed with phosphate-buffered saline (PBS) (Mediatech, Inc.; CORNING, Manassas, VA, USA) and lysed on ice for 1 h using RIPA lysis buffer (EMD Millipore, Billerica, MA, USA) supplemented with protease and phosphatase inhibitor cocktails (Cell Signaling Technology, Inc., Danvers, MA, USA). Lysates were centrifuged at 10,000 × g for 30 min at 4 °C, and the supernatants were collected. Protein concentrations were quantified using the Bradford assay (BioShop Canada Inc., Burlington, ON, Canada) with bovine serum albumin (BSA) (BioShop Canada Inc.) as the standard. Equal amounts of protein (20–40 μg) were mixed with 5X Laemmli sample buffer (ACE Biolabs, Taiwan), heated at 95 °C for 5 min, and separated by 10% SDS-PAGE. Proteins were transferred onto methanol-activated polyvinylidene difluoride (PVDF) membranes (EMD Millipore). Membranes were blocked with 5% BSA in Tris-buffered saline containing 20 mM Tris (Bioman Scientific Co., Ltd.), 150 mM NaCl (Shimakyu’s Pure Chemicals, Osaka, Japan), and 0.1% Tween-20 (Bioman Scientific Co., Ltd.) (TBST) for 1 h at room temperature, followed by overnight incubation at 4 °C with primary antibodies targeting HIF-1α (cat. no. 36169), poly (ADP-ribose) polymerase (PARP) (cat. no. 9542), MTH1 (cat. no. 43918), SOD2 (cat. no. 13141) (all from Cell Signaling Technology, Inc.), and GAPDH (cat. no. GTX100118; GeneTex, Inc., Irvine, CA, USA) as a loading control. Primary antibodies were applied at a dilution of 1:1000. After washing with TBST, membranes were incubated with horseradish peroxidase-conjugated secondary antibodies (cat. no. 111-035-003; Jackson ImmunoResearch Laboratories, Inc., West Grove, PA, USA) at a 1:10,000 dilution for 1 h at room temperature. Protein bands were detected using enhanced chemiluminescence (Advansta, San Jose, CA, USA) and imaged with the Amersham Imager 600 system (GE Healthcare, Chicago, IL, USA) for densitometric analysis.

### Analysis of ROS generation

Intracellular ROS levels were determined using FACSVerse flow cytometer (BD Biosciences, San Jose, CA, USA). Following Bai, TCS, or their combination treatment, A549 cells were harvested using trypsin-EDTA (Gibco; Thermo Fisher Scientific, Inc.), washed twice with PBS, and subsequently resuspended in 5 µM dihydroethidium (DHE) solution (Sigma-Aldrich) prepared in PBS. The samples were incubated in the dark at 37 °C for 30 min to facilitate dye uptake. Fluorescence intensity was then measured by flow cytometry, and the data were analyzed with FlowJo Version 10.0.0 software (FlowJo LLC, Ashland, OR, USA).

### Analysis of the MMP

MMP was evaluated through flow cytometric analysis. Following Bai, TCS, or their combination treatment, A549 cells were harvested using trypsin-EDTA, washed twice with PBS, and resuspended in 40 nM 3,3’-dihexyloxacarbocyanine iodide (DiOC_6_(3)) (Enzo Life Sciences, Farmingdale, NY, USA) prepared in PBS. The samples were incubated in the dark at 37 °C for 30 min to allow dye uptake. DiOC_6_(3) is a voltage-sensitive dye that preferentially accumulates in mitochondria with intact membrane potential, producing strong green fluorescence. A reduction in MMP diminished dye accumulation, thereby decreasing fluorescence intensity [43]. Fluorescence measurements were conducted via flow cytometry, and data were processed using FlowJo software.

### Apoptosis analysis

Apoptotic cell populations were assessed via flow cytometry. Following Bai, TCS, or their combination treatment, A549 cells were harvested using trypsin-EDTA, washed twice with PBS, and stained with Annexin V–FITC and propidium iodide (PI) (BD Biosciences). Annexin V binds to phosphatidylserine (PS), which is translocated to the outer leaflet of the plasma membrane during early apoptosis, resulting in green fluorescence [44]. PI is a nucleic acid-binding dye that penetrates cells with compromised membranes, staining late apoptotic or necrotic cells and emitting red fluorescence. For staining, Annexin V–FITC was diluted 1:500, and PI was applied at a final concentration of 500 ng/mL in Annexin V binding buffer. Samples were incubated in the dark at 37 °C for 30 min to allow dye uptake. Early and late apoptotic populations were quantified based on fluorescence signals, with data analyzed using FlowJo software.

### Statistical analysis

All experiments were conducted at least in triplicate. Data are presented as mean ± standard deviation (SD). Statistical analyses were performed using OriginPro 2015 Version 9.2 software (OriginLab Corporation, Northampton, MA, USA). Statistical differences between groups were assessed by one-way analysis of variance (ANOVA) followed by Tukey’s post hoc test. A *p*-value of less than 0.05 was considered statistically significant.

## Results

### Establishment of the hypoxic cell culture model

Given the crucial role of hypoxia plays in cancer pathophysiology, CoCl_2_, a widely used chemical hypoxia mimetic, was employed to induce hypoxic conditions *in vitro* [14]. This study aimed to evaluate the effects of Bai and TCS on A549 cells under hypoxic conditions. Based on previous research [45], a concentration of 150 μM CoCl_2_ was selected for subsequent experiments. To confirm that this concentration effectively induced hypoxia without causing significant cytotoxicity, an MTT assay was conducted after 48 h of CoCl_2_ treatment on A549 cells (Fig 2A). The results demonstrated that cell viability remained around 96% relative to untreated control, indicating negligible cytotoxic effects and no substantial decline in viability. Therefore, 150 μM CoCl_2_ was used throughout all *in vitro* experiments to establish hypoxia in A549 cells.

**Fig 2.**
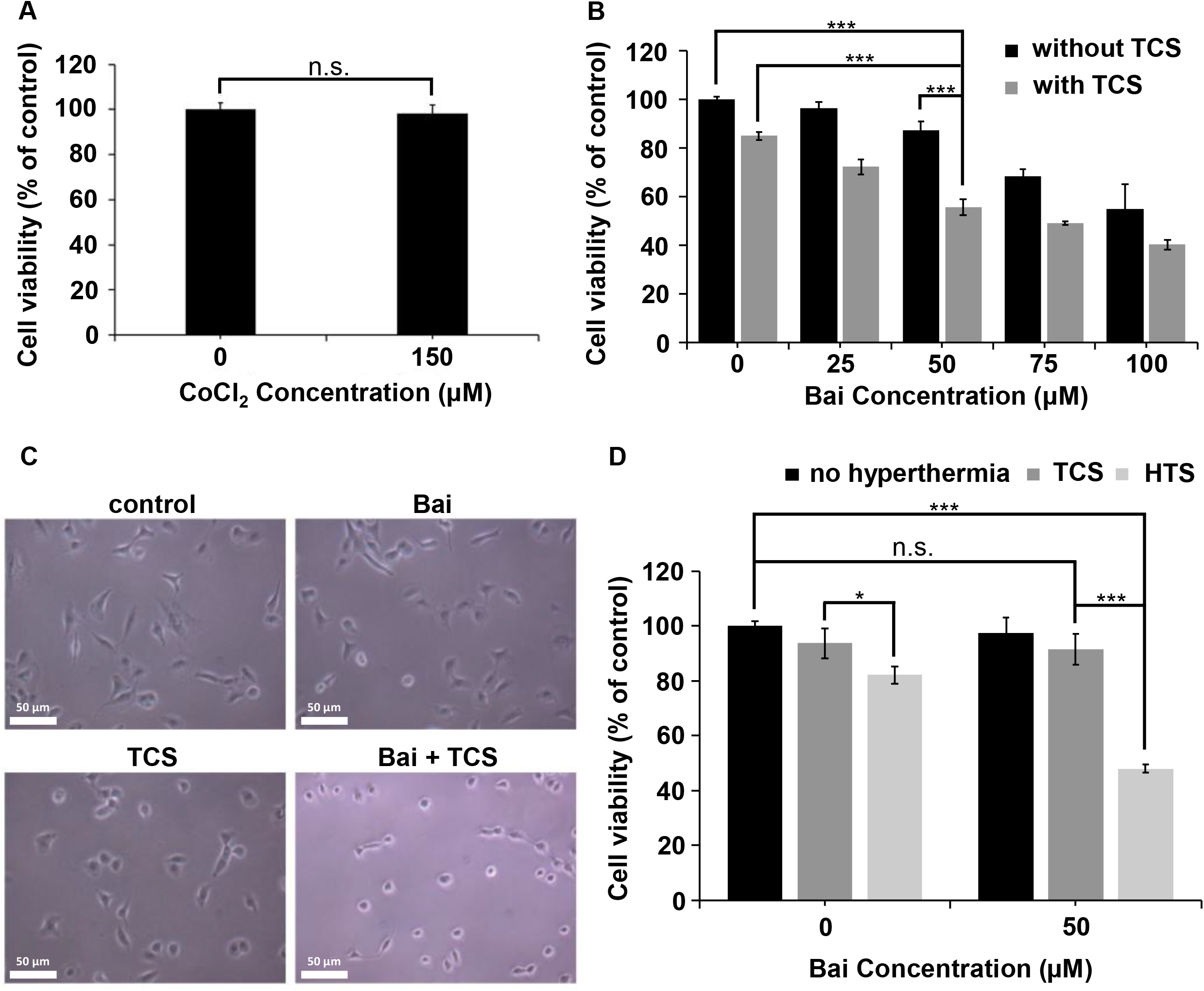
Assessment of cell viability in A549 and IMR-90 cells by MTT assay. (A) Effects of various CoCl_2_ concentrations (0, 30, 60, 90, 120, 150 µM) on the viability of A549 cells after 48 h of treatment. (B) Following a 6 h pre-treatment with 150 µM CoCl_2_ to induce hypoxia, A549 cells were exposed to different concentrations of Bai (0, 25, 50, 75, 100 µM) either alone or in combination with TCS. Cell viability was measured 48 h after treatment. (C) Morphological alterations in A549 cells were observed under control, Bai (50 µM), TCS, and combined Bai (50 µM) + TCS treatment. (D) IMR-90 cells were treated with or without 50 µM Bai, in combination with either TCS or HTS, and cell viability was measured after 48 h. Cell viability data were normalized to the control group and presented as mean ± SD from at least three independent experiments. Statistical significance between the indicated groups is indicated as *P < 0.05, **P < 0.01, and ***P < 0.001; non-significant results are denoted as n.s.

### Synergistic inhibition of A549 cell viability by Bai and TCS under hypoxic conditions

To explore the combined anticancer effects of Bai and TCS under CoCl_2_-induced hypoxia, the viability of A549 NSCLC cells was assessed using the MTT assay. Cells were initially exposed to CoCl_2_ for 6 h to induce hypoxia, followed by treatment with varying concentrations of Bai (25 μM, 50 μM, 75 μM, and 100 μM) [46]. Subsequently, cultures were treated with TCS to determine the optimal synergistic dosage, and cell viability was measured 48 h after treatment. As shown in Fig 2B, treatment with 50 μM Bai alone reduced cell viability to about 90% of control level, while TCS alone decreased viability to roughly 85%, indicating minimal cytotoxicity when administered individually. Notably, the combined treatment of Bai (50 μM) and TCS further lowered viability to approximately 60%, demonstrating a significant synergistic effect comparable to that observed with100 μM Bai alone (Fig 2B). The SQ value for Bai at 50 μM was 1.58, exceeding those at other concentrations and signifying the most pronounced synergistic interaction (Table 1). These findings suggest that co-treatment under hypoxic conditions synergistically inhibits A549 cell viability. Based on these results, 50 μM Bai combined with TCS was selected for further experimental analyses. Morphological analysis of A549 cells further supported these findings (Fig 2C). While treatment with either Bai or TCS alone induced only minimal morphological alterations, their combined treatment elicited pronounced changes in cellular morphology relative to the control group.

**Table 1.** SQ Value Analysis for Synergism.

| Bai concentration (μM) | 25 | 50 | 75 | 100 |
| --- | --- | --- | --- | --- |
| Inhibition rate of Bai (%) | 3.91 | 12.97 | 31.77 | 45.24 |
| Inhibition rate of TCS (%) | 15.23 | 15.23 | 15.23 | 15.23 |
| Inhibition rate of<br>Bai + TCS (%) | 27.92 | 44.56 | 51.02 | 59.97 |
| SQ value | 1.46 | 1.58 | 1.09 | 0.99 |

### Tolerance of normal lung cells to TCS treatment

To evaluate whether TCS mitigates damage to normal lung cells, the normal human lung fibroblast cell line IMR-90 was employed in the experiments. Cells were subjected to treatment with Bai, TCS, or their combination using the same procedure as with A549 cells, thereby enabling a direct comparison of the effects of TCS and conventional HTS on cell viability. As shown in Fig 2D, administration of 50 μM Bai in combination with TCS maintained cell viability at approximately 90% relative to the control group, with no statistically significant difference in IMR-90 cells. In contrast, treatment with the same concentration of Bai combined with conventional HTS on IMR-90 cells significantly reduced viability to about 50% of control, indicating considerable cellular damage induced by HTS. These results suggest that TCS substantially mitigates thermal injury in normal lung cells compared to traditional HTS.

### Reduction of HIF-1α expression by combined Bai and TCS treatment

To confirm the efficacy of CoCl_2_ in simulating hypoxic conditions, we first assessed the expression of HIF-1α, a protein stabilized under hypoxia and implicated in cancer progression [10, 12, 13]. Western blot analysis revealed that treatment of A549 cells with 150 μM CoCl_2_ resulted in a pronounced elevation of HIF-1α protein levels (Fig 3A), thereby validating this concentration as an effective *in vitro* model of hypoxia. It is known that HIF-1α functions as a pivotal transcription factor facilitating cellular adaptation and survival under hypoxic stress [13]. Its overexpression promotes tumor progression, whereas its inhibition has been proposed as a potential therapeutic approach [47]. Subsequently, the modulatory effects of Bai and TCS on HIF-1α expression were investigated in CoCl_2_-treated A549 cells. As shown in Fig 3B, treatment with either Bai or TCS alone reduced HIF-1α expression to approximately 80% of control level, whereas their combined application elicited a more substantial reduction to about 50%, indicating a synergistic inhibitory effect.

**Fig 3.**
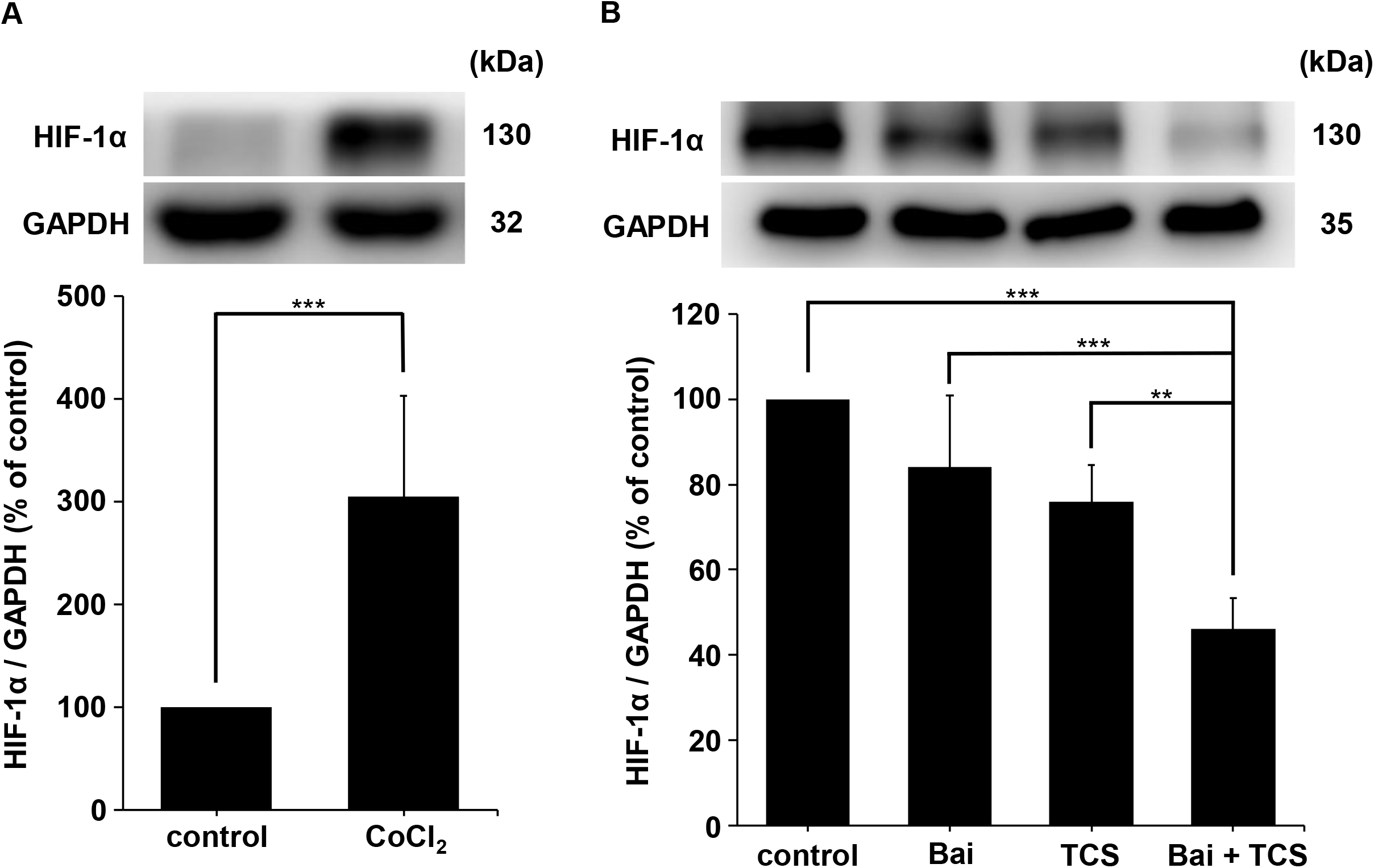
Expression of HIF-1α protein in A549 cells. (A) Western blot analysis of HIF-1α expression in A549 cells subjected to treatment with or without 150 µM CoCl_2_ to simulate hypoxic conditions. (B) Quantification of HIF-1α expression in CoCl_2_-treated A549 cells after treatment with Bai, TCS, and their combination. GAPDH was used as the loading control. Protein expression levels were normalized to the control group and presented as mean ± SD from at least three independent experiments. Statistical significance between the indicated groups is indicated as *P < 0.05, **P < 0.01, and ***P < 0.001.

### Bai combined with TCS increases intracellular ROS levels

Excessive accumulation of ROS can trigger mitochondria-mediated intrinsic apoptosis [48]. In cancer cells, HIF-1α plays a crucial role in regulating antioxidant defenses [15], rendering ROS modulation which is a key factor in cancer cell survival. As previously demonstrated, the combination of Bai and TCS notably decreased HIF-1α expression, which may weaken cellular antioxidant capacity. To assess the effect of this combined treatment on ROS production, intracellular superoxide levels were quantified via flow cytometry after staining with DHE, a common fluorescent probe for detecting superoxide radicals [49] (Figs 4A and 4B). Treatment with either Bai or TCS alone elevated ROS levels to approximately 120% of control, whereas their combination significantly increased ROS accumulation to roughly 150%, demonstrating a synergistic enhancement of oxidative stress. These findings suggest that co-treatment with Bai and TCS cooperatively enhances ROS production, which may lead to increased cellular damage and decreased viability.

**Fig 4.**
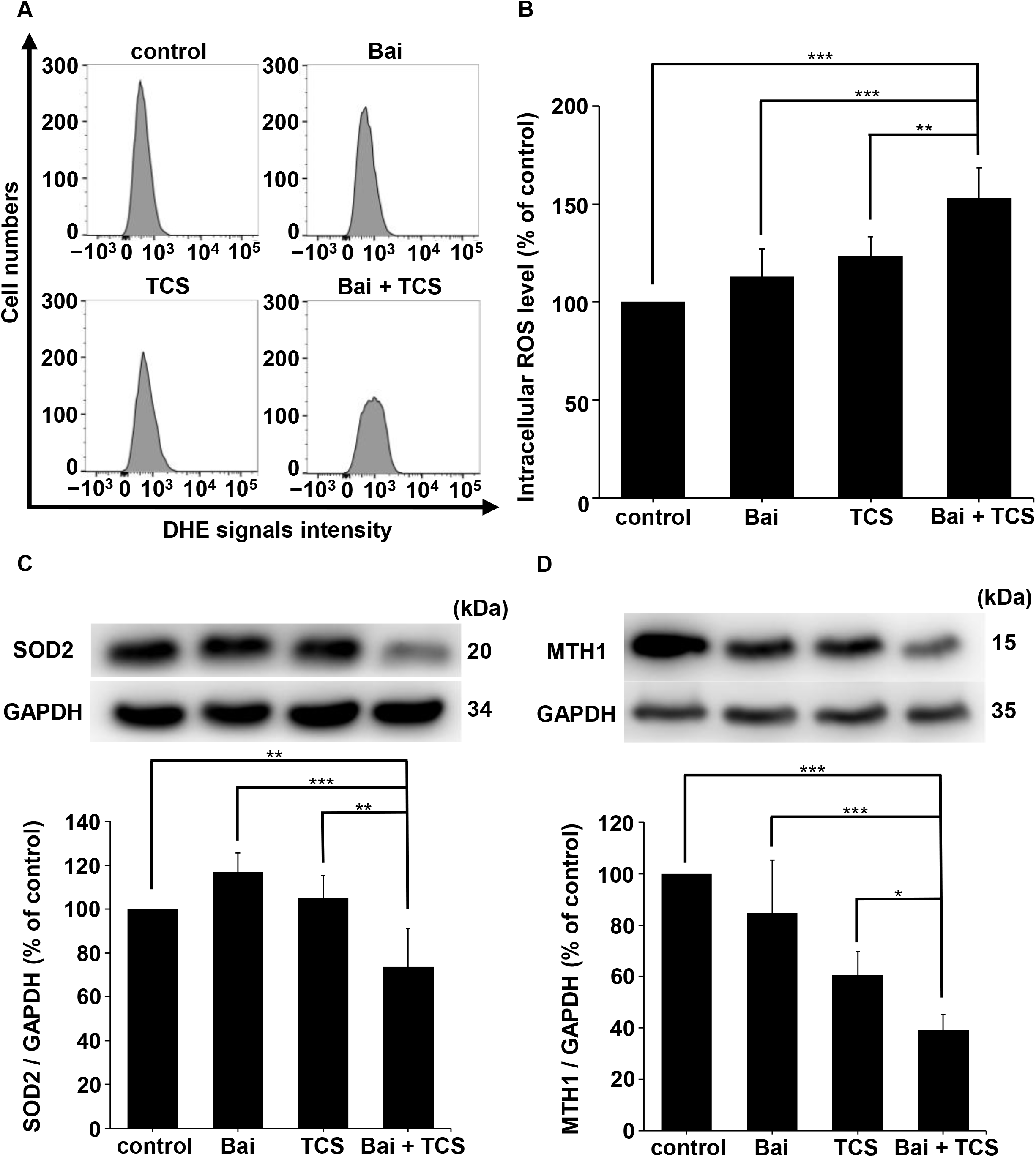
Oxidative stress-associated alterations in A549 cells subjected to hypoxic conditions. (A) Flow cytometry histograms showing the distribution of DHE fluorescence intensity distribution in A549 cells after treatment with Bai, TCS, or their combination. (B) Quantification analysis of the mean DHE fluorescence intensity for each treatment group. Western blot analysis showing the expression levels of (C) SOD2 and (D) MTH1 proteins, respectively, in A549 cells exposed to hypoxia following the indicated treatments. GAPDH was used as the loading control. All data were normalized to the control group and presented as mean ± SD from at least three independent experiments. Statistical significance between the indicated groups is indicated as *P < 0.05, **P < 0.01, and ***P < 0.001.

### Bai combined with TCS downregulates SOD2 expression

SOD2 is a critical mitochondrial antioxidant enzyme that catalyzes the dismutation of superoxide into the less reactive hydrogen peroxide [19], thus mitigating cellular damage induced by ROS. In this study, Western blot analysis was performed to assess SOD2 expression after treatment (Fig 4C). Neither Bai nor TCS alone elicited a significant alteration in SOD2 levels relative to the control group. However, the combined treatment produced a pronounced reduction, decreasing SOD2 protein expression to approximately 70% of the control. These findings suggest that the combination may suppress SOD2 expression via downregulation of HIF-1α, consequently weakening antioxidant defense mechanisms. As a result, the reduced capacity for ROS scavenging may contribute to ROS accumulation and enhanced cytotoxicity in A549 cells.

### The combination of Bai and TCS suppresses the expression of MTH1

Given the observed elevation in ROS levels caused by the combined Bai and TCS treatment, we further investigated the expression of MTH1. MTH1 is an enzyme that prevents the incorporation of oxidized nucleotides into DNA and RNA, thereby protecting cells from oxidative damage [50, 51]. Western blot results (Fig 4D) showed that the combination treatment significantly decreased MTH1 expression to roughly 40% of control level. This downregulation likely disrupts the nucleotide pool sanitization system, weakening the cellular capacity to defend against oxidative DNA damage. Consequently, the impaired DNA repair capacity may lead to more DNA lesions and promote the activation of apoptotic pathways.

### Synergistic effect of Bai and TCS in enhancing MMP disruption

Building upon our previous findings that the combination of Bai and TCS elevates ROS levels and suppresses MTH1 expression, this outcome would impose oxidative stress on the cells and reduce the cells’ capability to repair ROS-induced DNA damage [17, 50, 51], and we hypothesized that this treatment may also compromise mitochondrial integrity. To investigate this, MMP was assessed via flow cytometry (Figs 5A and 5B). The results indicated that approximately 25% of cells treated with the combined treatment exhibited reduced MMP, representing a significant increase compared to cells treated with either Bai or TCS treatment alone. These data demonstrate that the combined treatment synergistically disrupts mitochondrial function in A549 cells, which may contribute to the initiation of apoptosis.

**Fig 5.**
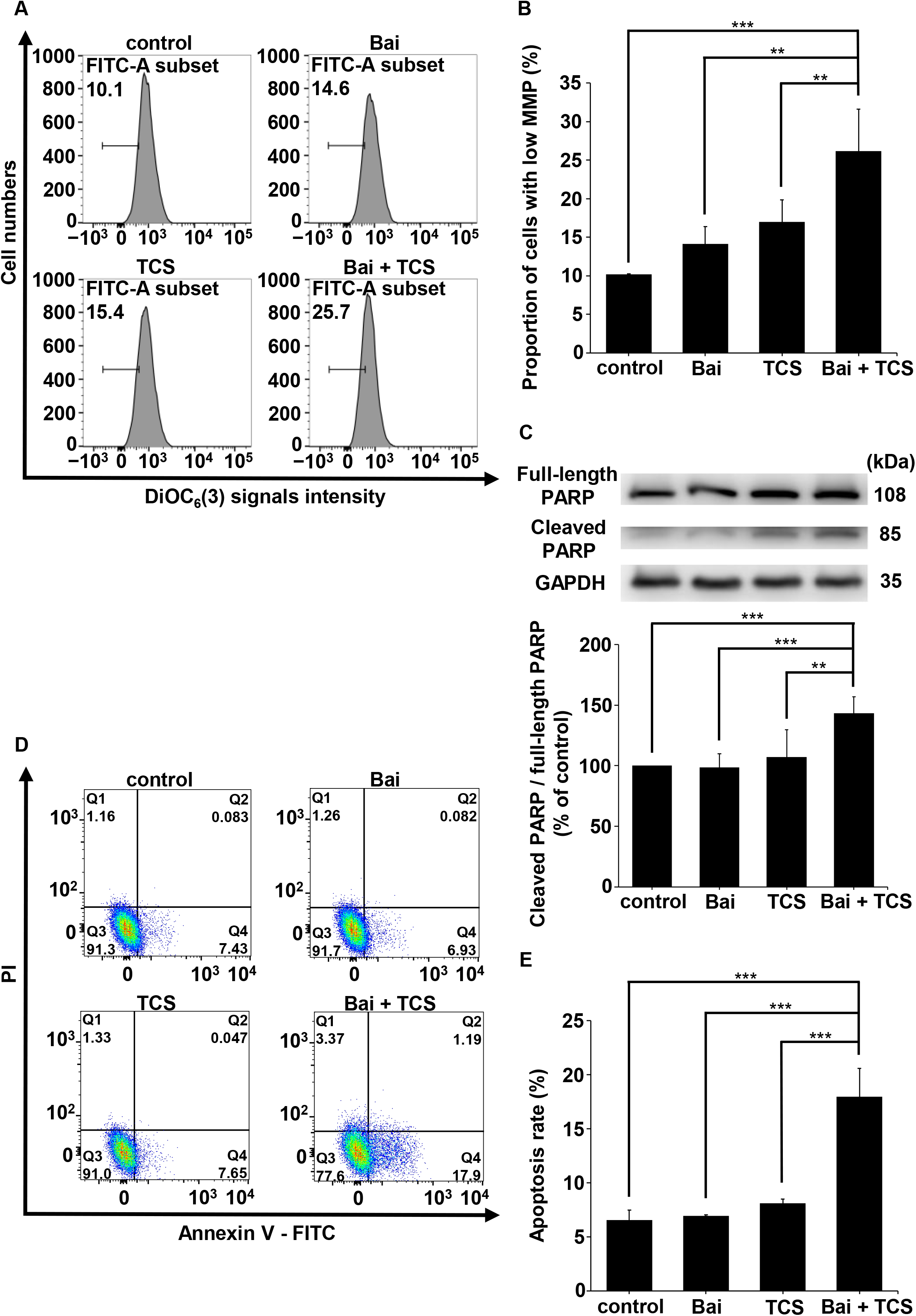
Induction of apoptosis in hypoxic A549 cells. (A) Flow cytometry histograms showing DiOC_6_(3) fluorescence intensity, indicative of MMP, following treatment with Bai, TCS, and their combination. (B) Quantitative analysis of the proportion of cells with reduced DiOC_6_(3) fluorescence intensity, corresponding to decreased MMP. (C) Western blot analysis of cleaved and full-length PARP was performed. The ratio of cleaved to full-length PARP was utilized as a marker of apoptosis, with GAPDH serving as a loading control. (D) Flow cytometric analysis of apoptosis in CoCl_2_-treated A549 cells using Annexin V-FITC and PI staining. (E) Quantification of total apoptotic cells, calculated as the sum of early apoptotic (Q3) and late apoptotic (Q2) cells shown in (D). All data were normalized to the control group and presented as mean ± SD from at least three independent experiments. Statistical significance between the indicated groups is indicated as *P < 0.05, **P < 0.01, and ***P < 0.001.

### Bai and TCS synergistically induce apoptosis in A549 cells

PARP cleavage serves as an indicator of DNA repair inhibition and apoptosis, occurring downstream of MMP loss and caspase activation [52, 53]. As shown in Fig 5C, Western blot analysis demonstrated that the combined administration of Bai and TCS significantly elevated cleaved PARP levels to approximately 140% relative to the control, whereas individual treatments resulted in only slight increases (100–110%), consistent with the observed reduction in MMP. Along with previous findings of increased ROS and reduced MTH1 expression, these results suggest that the combined treatment causes mitochondrial dysfunction and initiates apoptotic processes. Apoptosis progresses through distinct phases: early apoptosis is characterized by the externalization of PS on the cell membrane, which is specifically recognized by Annexin V, whereas late apoptosis involves loss of membrane integrity, allowing PI to penetrate and bind DNA [44, 54]. Flow cytometry analysis (Fig 5D) differentiated live cells (Q4), early apoptotic cells (Q3), late apoptotic cells (Q2), and necrotic cells (Q1). The combined treatment significantly increased the total apoptotic cell population (early plus late apoptosis: control 6%, Bai 7%, TCS 8%, and Bai + TCS 18%) compared to single treatments and control (Fig 5E), thereby confirming an enhanced induction of apoptosis.

## Discussion

This study aims to explore the combined anticancer effects of Bai and TCS on A549 NSCLC cells under hypoxic conditions. Tumor environments often experience hypoxia [6, 7], which leads to the stabilization and accumulation of HIFs, particularly HIF-1α [7, 10]. As a crucial regulator for cellular adaptation to oxygen deficiency, HIF-1α modulates the expression of genes involved in angiogenesis, metabolic reprogramming, and survival pathways. Persistent activation of HIF-1α under hypoxic conditions promotes tumor growth and aggressiveness, making it as a critical mediator in cancer progression [10–13]. In this study, CoCl_2_ was utilized as a chemical hypoxia-mimetic agent that stabilizes HIF-1α by inhibiting its degradation, thereby simulating hypoxic conditions [14]. Treatment with 150 µM CoCl_2_ did not harm A549 cell viability, as shown by MTT assays (Fig 2A), and Western blot analysis confirmed elevated HIF-1α levels, thereby validating the hypoxia model for subsequent experiments (Fig 3A).

Previous studies have shown that Bai exhibits multiple anticancer activities and suppresses HIF-1α expression in various cancer models [21–24]. However, its role in modulating hypoxia-induced HIF-1α expression in NSCLC cells has not been fully characterized. Furthermore, the therapeutic potential of Bai is limited by its poor aqueous solubility and low bioavailability [25–27], which may restrict its effectiveness in the tumor microenvironment. Given that hypoxia is a critical feature of solid tumors [6, 7], developing strategies to enhance Bai-mediated inhibition of hypoxia-associated signaling pathways is highly desirable. Therefore, a combination strategy incorporating TCS was employed to enhance the anticancer efficacy of Bai. The treatment involved subjecting cells to 10 cycles of heating at about 43°C for 3 min followed by cooling to around 41°C for 30 s, a protocol previously reported to maximize therapeutic benefits while minimizing thermal injury to normal cells [34, 36, 37]. MTT assay results indicated that the combination of 50 µM Bai with TCS synergistically decreased A549 cell viability, producing effects comparable to those observed with 100 µM Bai alone (Fig 2B). Additionally, morphological analysis revealed pronounced alterations in cellular morphology following the combination treatment on A549 cells (Fig 2C). Notably, this combined treatment did not cause cytotoxicity in IMR-90 normal human lung fibroblast cells, suggesting selective toxicity toward malignant cells (Fig 2D).

At the molecular level, prior investigations have demonstrated that conventional HTS can upregulate HIF-1α expression in A549 cells under specific conditions [55, 56]. In our experimental system, however, treatment with TCS resulted in a reduction of HIF-1α levels (Fig 3B), indicating that the regulation of HIF-1α may be highly contingent upon the particular thermal parameters employed in different hyperthermia modalities. Given that HIF-1α modulates antioxidant defense pathways [15, 18], the suppression of HIF-1α induced by TCS could explain the observed increase in ROS accumulation. Excessive ROS generation can deplete intracellular antioxidants, disrupt redox homeostasis, and inflict oxidative damage on cellular constituents [17]. Considering that cancer cells generally maintain elevated basal ROS levels relative to normal cells, strategies that further increase ROS or impair antioxidant defenses have been proposed as promising anticancer approaches, as they selectively surpass the oxidative stress tolerance of malignant cells [57]. Supporting this notion, previous studies have reported that both Bai and thermal stress independently elevate intracellular ROS levels in A549 cells [24, 31], which promotes apoptosis in cancer cells. Additionally, TCS has been shown to enhance ROS accumulation in PANC-1 human pancreatic cancer cells, thereby contributing to oxidative stress–mediated cytotoxicity [36]. In this study, we further demonstrated that the combined application of Bai and TCS significantly elevated intracellular superoxide levels in A549 cells compared to either treatment alone, indicating increased oxidative stress in the combination treatment (Figs 4A and 4B).

To elucidate the mechanism underlying this enhanced oxidative stress, we investigated the expression of SOD2, a key mitochondrial antioxidant enzyme responsible for catalyzing the dismutation of superoxide radicals into hydrogen peroxide [19]. Suppression of SOD2 has been reported to impair the antioxidant defenses of cancer cells, thereby diminishing their survival under oxidative stress conditions [58]. Importantly, HIF-1α has been identified as a transcriptional activator of the SOD2 gene, enhancing cellular resistance to hypoxia-induced oxidative stress [18]. Given our earlier observation that the combination treatment significantly lowered HIF-1α expression, we next explored whether SOD2 levels were also affected in A549 NSCLC cells. Our results showed that the combined treatment with Bai and TCS significantly suppressed SOD2 expression (Fig 4C), suggesting that compromised antioxidant defenses contributed to ROS accumulation and increased cytotoxicity. Moreover, excessive ROS can inflict damage on proteins, lipids, and nucleic acids, including oxidative modifications to DNA. To mitigate such damage, cells use defense mechanisms like MTH1, a DNA repair enzyme that removes oxidized nucleotides, to protect cells from ROS-induced DNA lesions during oxidative stress [50, 51]. Accordingly, we evaluated MTH1 expression in A549 cells treated with the combination of Bai and TCS. Previous research has shown that TCS alone can decrease MTH1 expression in A549 cells, thereby impairing the cells’ ability to repair oxidative DNA damage [37]. In our study, the combination of Bai and TCS further significantly lowered MTH1 expression (Fig 4D), indicating an additive or synergistic effect in weakening DNA repair processes. This reduction in MTH1 levels may exacerbate ROS-induced genomic instability, thereby contributing to the enhanced cytotoxicity observed with the combination treatment.

In addition to DNA damage, persistent oxidative stress detrimentally affects mitochondrial integrity and function [59]. Such mitochondrial dysfunction is recognized to activate the intrinsic apoptotic pathway [52, 53]. Given the crucial role of mitochondria in apoptosis, MMP was assessed as a marker of mitochondrial integrity and cellular homeostasis [60]. Previous research has shown that Bai alone can impair mitochondrial function and decrease MMP in A549 cells, thereby facilitating apoptosis [24]. To investigate the combined effects of Bai and TCS on mitochondrial function, MMP was measured via flow cytometry using DiOC_6_(3) staining [43]. The results revealed a significantly greater proportion of cells with reduced MMP in the combination treatment group compared to either Bai or TCS treatment alone (Figs 5A and 5B). The loss of MMP is known to initiate the caspase cascade, particularly caspase-3, which cleaves PARP—a multifunctional enzyme involved in DNA repair and programmed cell death—into 85 kDa and 24 kDa fragments, thereby inactivating its enzymatic activity [52]. Previous research has indicated that Bai and thermal stress can independently induce PARP cleavage in A549 cells, implicating their individual roles in apoptosis via mitochondrial dysfunction [24, 37].Consistent with these findings, our data demonstrated that while Bai or TCS alone caused a moderate increase in PARP cleavage (Fig 5C), their combination led to a significantly stronger effect, aligning with the observed mitochondrial impairment (Figs 5A and 5B). Furthermore, these effects were accompanied by a significantly elevated proportion of apoptotic cells, underscoring the enhanced apoptotic induction by the combined treatment (Figs 5D and 5E).

In summary, our study demonstrates that the combined administration of Bai and TCS synergistically diminishes the viability of A549 NSCLC cells without inducing excessive cytotoxicity in IMR-90 normal human lung fibroblast cells. This combination treatment notably suppresses HIF-1α expression under hypoxic conditions, which may reduce the transcriptional activation of the antioxidant enzyme SOD2, thereby elevating intracellular ROS levels and exacerbating oxidative stress. Concurrently, downregulation of MTH1 further impairs the cells’ capacity to repair ROS-induced DNA damage, thereby amplifying cytotoxic effects. These effects coincide with decreased MMP and increased PARP cleavage, indicative of mitochondrial apoptosis, which was confirmed by a significant increase in apoptotic cell populations following the combined treatment. Overall, these findings provide insights into the molecular mechanisms underlying the synergistic anticancer effects of Bai and TCS in CoCl_2_-induced hypoxic A549 cells. The combination effectively disrupts hypoxia-adaptive responses, intensifies oxidative stress, and promotes mitochondrial apoptosis. Taken together, these results suggest that combining Bai with TCS holds promising therapeutic potential for treating NSCLC in hypoxia-mimicking environments and merits further investigation.

## Data Availability Statement

All relevant data are within the manuscript and its supporting information files.

## Acknowledgments

The authors would like to acknowledge the service provided by the Research Core Facilities 3 Laboratory of Department of Medical Research at National Taiwan University hospital for use of flow cytometry system.

## Author Contributions

**Conceptualization:** Chih-Yu Chao.

**Data curation:** Yu-Wei Wang, Chih-Yu Chao

**Formal analysis:** Yu-Wei Wang, Chih-Yu Chao

**Methodology:** Yu-Wei Wang, Guan-Bo Lin, Yu-Yi Kuo, Chih-Yu Chao

**Investigation:** Yu-Wei Wang, Chih-Yu Chao

**Validation:** Yu-Wei Wang, Guan-Bo Lin, Fang-Tzu Hsu, Yu-Yi Kuo, Yun-Hao Chen, Chih-Yu Chao

**Funding asquisition:** Chih-Yu Chao

**Supervision:** Chih-Yu Chao

**Writing-original draft:** Yu-Wei Wang, Chih-Yu Chao

**Writing-review & editing:** Chih-Yu Chao

## Abbreviations

ANOVA: analysis of variance
Bai: baicalein
BSA: bovine serum albumin
CoCl_2_: cobalt (II) chloride
ddH_2_O: deionized and distilled water
DHE: dihydroethidium
DiOC_6_(3): 3,3’-dihexyloxacarbocyanine iodide
DMEM: Dulbecco’s modified Eagle’s medium
DMSO: dimethyl sulfoxide
HIFs: hypoxia-inducible factors
HIF-1: hypoxia-inducible factor-1
HIF-1α: hypoxia-inducible factor-1 alpha
HTS: hyperthermia stimulation
H_2_O_2_: hydrogen peroxide
TCS: thermal-cycling stimulation
MMP: mitochondrial membrane potential
MTH1: MutT homolog 1
MTT: 3-(4,5-dimethylthiazol-2-yl)-2,5-diphenyltetrazolium bromide
NSCLC: non-small cell lung cancer
PARP: poly (ADP-ribose) polymerase
PBS: phosphate-buffered saline
PI: propidium iodide
PS: phosphatidylserine
PVDF: polyvinylidene difluoride
SD: standard deviation
SDS: sodium dodecyl sulfate
SOD2: superoxide dismutase 2
SQ: synergism quotient
ROS: reactive oxygen species

